# Phylogenomics of *Messor* harvester ants (Hymenoptera: Formicidae: Stenammini) unravels their biogeographical origin and diversification patterns

**DOI:** 10.1101/2025.02.13.638115

**Authors:** Yannick Juvé, Arthur Weyna, Elodie Lauroua, Sabine Nidelet, Mourad Khaldi, Ghania Barech, Claude Lebas, Jean-Yves Rasplus, Astrid Cruaud, Fabien L. Condamine, Jonathan Romiguier

## Abstract

As a major abiotic factor, climate change is expected to profoundly alter biological communities. On this basis, identifying how past temperature variations affected species diversification and distribution can help to predict the effects of the ongoing climate change. In this study, we focused on the harvester ant genus *Messor* which is mostly adapted to dry environments dominated by seed-producing plants. The phylogenetic analysis of 2,524 markers obtained from 58 *Messor* species/subspecies, supports their emergence in the Irano-Indian area approximately 20 Mya. Phylogenetic relationships uncovered in this study enabled us to redefine historical taxonomic groups, providing a solid basis for future revisions that encompass the entire genus diversity. Their diversification appears to be affected by temperature, with a higher speciation rate during warmer periods. This confirms that the ecological specialization of *Messor* makes them strongly dependent on thermal conditions. Our results highlight the importance of abiotic factors on diversification processes, especially for highly specialized species that may exhibit predictable evolutionary responses to climate changes.

## I. Introduction

Climate change is expected to influence species distributions in complex ways by influencing both the biotic and abiotic components of ecosystems (Bellard *et al*., 2012). This interplay challenges our ability to predict distribution shifts or local extinction of species (Parmesan *et al*., 1999; Urban *et al*., 2016). However, not all species are equal in this regard, and some species can be expected to respond in more predictable ways than others to climate change. This is the case of biome specialists who should be strongly impacted (Vrba, 1980, 1987), while biome generalists should be less dependent on abiotic factors (Gamboa *et al*., 2022).

Based on climate and vegetation zonation, biome distribution is heterogeneous at the global scale with for instance the Palearctic region being composed of at least six types of biome (Walter, 1970; Donoghue and Edwards, 2014). However, some of these biomes expanded or disappeared over time (Pound *et al*., 2012; Pound and Salzmann, 2017). During the Cenozoic, when the climate cooled down (Westerhold *et al*., 2020), the tropical biomes vanished or shifted southward (Ziegler *et al*., 2003; Pound *et al*., 2012). These changes impacted diversity and diversification at global (Meseguer and Condamine, 2020; Quintero *et al*., 2023) and regional (Graham, 2011; Jaramillo, 2023) scales. During the Neogene, and more specifically during the last 11 million years (Westerhold *et al*., 2020), global climate cooling induced substantial aridification over the Northern Hemisphere (Eronen *et al*., 2012). One of the main biome shifts over the late Cenozoic consisted of the expansion of grasslands (Edwards *et al*., 2010; Strömberg, 2011; Pound *et al*., 2012; Spriggs, Christin and Edwards, 2014). Most grassland ecosystems are dominated by grasses (family Poaceae) that employ C_4_ photosynthetic pathways, which are better adapted to dry and low CO_2_ conditions (Strömberg, 2011). Although their origin remains debated, fossil and phylogenetic data indicate an early Miocene diversification and a Pliocene dominance of grasses linked with the drop in atmospheric CO_2_ (Palazzesi *et al*., 2022).

The combination of climate and biome changes is thought to have played a key role in the evolution of species communities, notably by promoting the diversification of phytophagous lineages associated with grasslands (e.g. Kergoat *et al*., 2018; Liu, Fu and Luo, 2024). There is growing evidence that past climate change has influenced the pace at which groups of organisms diversified (Benton, 2009; Erwin, 2009). Analyses of the fossil record and/or dated phylogenies suggest that past temperature fluctuations could likely be a driver of speciation rates in many clades (e.g. Condamine, Romieu and Guinot, 2019). The general consensus is that speciation rates were higher when the climate was warmer, which led to a slowdown of speciation toward the present (Condamine, Rolland and Morlon, 2019). However, we still lack macroevolutionary studies on invertebrates and insects supporting this consensus (but see (Condamine *et al*., 2015, 2018; Kergoat *et al*., 2018; Baird *et al*., 2021).

Within insects, the harvester ant genus *Messor* Forel, could represent a relevant biological model to better assess the role of climate change in insect diversification. It is highly specialized for granivory (Lévieux and Diomande, 1978; Hahn and Maschwitz, 1985), and is mainly found in open areas of semi-arid to arid habitats of the Old World. Their interaction with seed-producing plants makes them abundant around the Mediterranean basin, but also extending to South Africa and Japan (Plowes, Johnson and Hölldobler, 2013). Within the subfamily Myrmicinae, *Messor* currently includes 134 species (based on the AntCat database, antcat.org, accessed on 13 November 2024), making it the second most diverse genus in the tribe Stenammini (7 genera, 458 species). Based on our current knowledge, two distant regions (Western North Africa and Middle-East) comprise a large part of the species diversity. According to AntMaps (Guenard *et al*., 2017), 30 species of *Messor* are recorded from Morocco and Iran. The emergence of the genus has been previously inferred to be within the Palearctic, between 12.6 and 16.13 Mya (Branstetter *et al*., 2022), however with only a few species included in the analysis. Whether the biogeographic origin of *Messor* corresponds to one of the current diversity hotspots, or whether this diversity pattern is not representative of their ancestral area, remains an open question. It is also unknown if the highly specialized ecology of harvester ants conditioned their diversification. Because aridification and habitat opening promoted the dominance of seed-grass plants (Hui *et al*., 2021), one can wonder whether such environmental transitions also shaped the diversification of the granivorous *Messor* ants. As a genus mostly associated with arid biomes, we thus hypothesize that its diversification patterns and biogeographical history could have been strongly determined by past climatic changes.

Several challenges may hamper our ability to unravel the origin and diversification of the genus *Messor*. In particular, most of the species are difficult to identify and classify. Because of the continuous worker polymorphism which strongly distorts allometry (Schlick-Steiner *et al*., 2006), accurate descriptions are scarce and identification mostly depends on specimen size (Salata and Borowiec, 2019). Large individuals with large heads (majors) are preferred as they present more marked morphological features (Bernard, 1981), but without proper sampling or colony maturity, species identification is difficult if not impossible when examining smaller workers. Sexual caste would help to better grasp and categorize their diversity, but their observation is mostly seasonal-dependent and many of them are still unknown or undescribed (Bernard, 1953). Species of *Messor* can also exhibit strong variations between populations of what is presumably the same species (Bernard, 1955).On the contrary, entities that are supposed to represent distinct species are morphologically extremely close, not to say cryptic (Bernard, 1985; Steiner *et al*., 2018). In addition, most of the morphological characters used to separate species appear homoplasic, making species characterization difficult. Third, even if only a few taxa have their types lost or destroyed, many are only represented in a single museum, which makes it even more complicated to assess the validity of enigmatic or dubious taxa that often lack precise or complete descriptions. Considering their widespread distribution and their arduous diagnosis, regional taxonomic studies might have promoted taxon synonymy. Some studies have been conducted on broad geographic scales and taxonomic ranges, such as in the Maghreb (Barech *et al*., 2020) or the Afrotropics (Bolton, 1982), but none has yet tried to investigate and link most of the known diversity in a global taxonomic assessment. Finally, some species appear to hybridize (*M. minor* x *M. wasmanni*, Steiner *et al*., 2011; *M. mcarthuri* x *M. hellenius*, Lapeva-Gjonova and Borowiec, 2022) and others are known to possess complex reproductive systems called social hybridogenesis (in *M. barbarus*, *M. ebeninus* and *M. ibericus*; Romiguier *et al*., 2017; Steiner *et al*., 2018). If these phenomena frequently occur across the genus, they could be one of the main factors explaining morphological variability, and thus the difficulty in identifying consistent species boundaries.

Facing these difficulties, it appears challenging but nonetheless essential to complement morphology-based systematics with genomics to clarify the higher-level systematics and species-group taxonomy of the genus *Messor*. Here, we aimed to infer a dated species-level phylogeny of *Messor* using genomic data to lay the systematic foundation of the genus, and study their evolutionary history by estimating their most likely ancestral origin and dispersal routes to explain the current diversity pattern. We also explored the deep-time environmental factors tied to their diversification. A first attempt has been conducted in a phylogenetic study of the tribe Stenammini, but this study contained only six sequenced *Messor* species with limited geographic representation (Branstetter *et al*., 2022). To fill this gap, we sequenced 58 *Messor* species/subspecies from 23 countries across the genus geographic range and reconstructed their evolutionary history through a phylogenomic approach including molecular dating and macroevolutionary analyses. We paid attention to identify potential cases of hybridization, which could create discordant gene topologies and low branch supports.

## II. Material and methods

### 1. Taxon sampling and sequencing

Among the investigated specimens, 62 samples were obtained from the ant collection hosted at the London British Museum of Natural History (BMNH) in 2019, and 31 were sampled between 2005 and 2023 in alcohol then stored in the University of Montpellier freezers (Table S1). Non-destructive extraction was achieved using the Qiagen (Valencia, CA) DNeasy Blood and Tissue kit, following the manufacturer’s protocol with a few modifications, as detailed in (Cruaud et al., 2019) and summarized hereafter. Extractions have been performed using the QIAGEN (Valencia, CA) DNeasy Blood & tissue kit, using a non-destructive method following the standard protocol with a few modifications, as detailed in Cruaud et al. (2019). To maximize the recovery of DNA and preserve the integrity of the specimens, small holes have been perforated with minutens in the three tagmata, mostly through integument membranes. Specimens were incubated overnight in an Eppendorf thermomixer (temperature = 56 C, mixing frequency = 300 rpm). Following incubation, specimens were removed, dried, and re-mounted using standard water-soluble and non-toxic glue. The remaining buffer was treated with ethanol to precipitate DNA and filtered using binding columns. To increase DNA yield, two successive elutions (50 L each) were performed with a heated AE buffer (56 C) and an incubation step of 15 minutes followed by centrifugation (8000 rpm for 1 minute at room temperature). Eppendorf microtubes LoBind 1,5ml were used for elution and stored at -20 C until library preparation. DNA was quantified with a Qubit 2.0 Fluorometer (Invitrogen).

To avoid sequencing contaminants inherent to conservation issues on dry specimens (mold/environmental DNA), we sequenced museum specimens after hybrid capture of 2,524 ant specific Ultra-Conserved Elements (UCE probes designed by Branstetter *et al*., 2017). UCE sequences are indeed adequate for fine taxonomical studies at species level, and still effective with low-quality samples from museums (Blaimer *et al*., 2016; Harvey *et al*., 2016) while being good markers for generating phylogenies, investigating complex species boundaries, and estimating divergence times (Blair *et al*., 2019). Sequencing libraries were prepared according to the protocol outlined by Faircloth *et al*. (2012, 2015) for Ultra-Conserved Elements capture, with modifications detailed in Cruaud *et al*. (2019). DNA was fragmented to a size of 400 bp using the Bioruptor Pico (Diagenode). Subsequent steps included end repair, 3’-end adenylation, adapter ligation, and PCR enrichment, all performed using the NEBNext Ultra II DNA Library prep kit for Illumina (NEB). Barcoded adapter pairs were utilized, containing amplification and Illumina sequencing primer sites, along with a 5 or 6 bp nucleotide barcode for sample identification. Pools of 16 samples were created at an equimolar ratio. UCE capture was accomplished by enriching each pool using the “Insect Hymenoptera 2.5K version 2, Ant-Specific” probe set (Branstetter et al., 2017) and MYbaits kits (MYcroarray, Inc.). The manufacturer’s protocol (MYbaits, user manual version 3) was followed. The hybridization reaction was conducted for 24 hours at 65°C. Post-enrichment amplification was performed on beads using the KAPA Hifi HotStart ReadyMix. Enriched libraries were quantified with Qubit, an Agilent Bioanalyzer, and qPCR using the Library Quantification Kit Illumina/Universal from KAPA (KK4824). The libraries were then pooled at an equimolar ratio. Paired-end sequencing (2x300bp) was carried out on an Illumina MiSeq platform at UMR AGAP (Montpellier, France).

For well-conserved specimens stored in alcohol, we sequenced whole genomes (WGS) as it allows more complete genomic analyses. DNA extractions have been done on the whole body of each sample using the Macherey-Nagel NucleoMag Tissue kit. Library preparation for whole genome sequencing has been performed using a custom Illumina protocol modified from Meyer and Kircher (2010) and detailed in Tilak *et al*. (2015).

To complement the taxonomic range of the sequenced specimens, especially for other Stenammini genera, UCE-capture data from Branstetter *et al*. (2022) has been added (see Table S1 for details). In total, the dataset of this study contains sequences of 192 specimens from 143 taxa investigated (51 sp. and 7 ssp. of *Messor* ; 29 sp. of *Aphaenogaster* ; 37 sp. of *Stenamma* ; 5 sp. of *Goniomma* ; 2 sp. of *Oxyopomyrmex* ; 9 sp. of *Veromessor* ; 3 sp. of *Novomessor*).

### 2. Phylogenomic inferences

We used phyluce v.1.7.1 (Faircloth, 2016) to identify UCE loci among 2,524 Formicidae UCEs references (probes designed by Branstetter *et al*., 2017). Each UCE locus were aligned using MAFFT v.7.490 (Katoh, Rozewicki and Yamada, 2019). We used trimaAl v1.2 (Capella-Gutiérrez, Silla-Martínez and Gabaldón, 2009) to only keep sites present in at least 50% of the sequences analyzed, then concatenated in a single supermatrix the trimmed sequences using catsequences v.1.5 (Creevey, Weeks and Ting, 2023). Phylogenetic analyses on the supermatrix obtained have been performed using IQ-TREE v.2.0.7 (Minh *et al*., 2020), setting up the substitution model GTR+F+I+G4 and 1,000 ultrafast bootstrap replicates (Hoang *et al*., 2018).

To produce gene trees, individual alignments for each locus have been trimmed with the heuristic method “-automated1” of trimAl v.1.2, and gene trees inferred with IQ-TREE (following the same options as above). These gene trees have been used to assess the gene and site concordance factors (gCF and sCF) of the supermatrix phylogeny (Minh *et al*., 2020). We then compared this supermatrix phylogeny with a tree-reconciliation approach using wASTRAL-hybrid (Zhang and Mirarab, 2022) to weigh the inference depending on the branch lengths and branch supports of every gene tree.

### 3. Detecting hybrids

Before investigating phylogenetics, genome heterozygosity has been estimated using a pipeline designed to detect first-generation hybrids (F1) in single individuals (Weyna *et al*., 2022). The underlying method partitions observed individual heterozygosity into two components: the component γ due to putative divergence between the two parental lineages, and the component θ due to polymorphism in the ancestral population of these parental lineages. Individuals with a significantly positive γ/θ ratio (i.e., with positive divergence between their parental lineages) can be regarded as F1 candidates. Here, we only considered ratios above 0.557, which corresponds to the lowest γ/θ value known so far among hybridogenetic species (*Paratrechina longicornis*, Weyna *et al*., 2022 supplementary material). Values of θ et γ < 0.0005 were considered as too low to be considered, regardless of the corresponding ratio value.

Both UCEs and BUSCOs (Benchmarking Universal Single-Copy Orthologs) can be used in estimating the heterozygosity, but only WGS specimens could be tested with the two genetic markers. Contigs of interest have been detected using phyluce v.1.7.1 with a 2,524 Formicidae UCE loci probe set (same as for phylogeny); and BUSCO v.5.3.1 with a 5,991 BUSCO dataset (Manni *et al*., 2021, dataset hymenoptera_odb10 from orthodb.org) and with *Camponotus floridanus* as the reference species for the integrated gene finder tool Augustus (Stanke *et al*., 2008). Per-contig counts of heterozygous sites have been performed with the snp-calling tool freebayes v.1.3.2 (Garrison and Marth, 2012), with an additional contamination filter. This process uses REF (allele of reference) and ALT (any other allele at REF locus) allelic depth to compute and compare genotype probabilities for each site. If the likelihood that the site is truly heterozygous is lower than the probability that the site is truly homozygous, then the SNP is rejected (see https://github.com/arthurWeyna/hybrid_scan for details). Because museomics are highly prone to contamination, we used a stringent value of 0.2 for the parameter e, which defines the expected frequency of ALT at true homozygous sites.

### 4. Estimating divergence times

To infer the divergence times of *Messor*, we relied on Bayesian relaxed clock inferences as implemented in MCMCtree v.4.10.7 (Yang, 2007) using the species tree obtained from the supermatrix analysis of the Stenammini. All *Messor* individuals have been included, but only one representative species for the other genera was retained (i.e. the one with the highest number of UCE loci). We did not use fossil calibration for divergence time estimation but we rather used secondary calibrations, with 95% height posterior densities (HPD) inferred by Branstetter *et al*., (2022). We retrieved the minimum and maximum ages of the 95% HPD for setting up uniform prior to calibrate five nodes (see Table S4) on a pruned Stenammini phylogeny, which besides *Messor* was represented by the best-covered specimen for other genera. To date the Stenammini, Branstetter *et al*., (2022) used the oldest known Stenammini fossil, †*Aphaenogaster dlusskyana* (Radchenko and Perkovsky, 2016), and a secondary calibration for the crown of Myrmicinae based on the phylogenetic study of Ward *et al*., (2015). We did not take into account the only known *Messor* species fossil, †*M. sculpturatus* (Carpenter, 1930), which has been attributed to the genus based on the presence of two closed cubital cells on the forewing; a feature that has since been identified as unreliable (Bolton, 1982). The molecular dataset was generated with the SortaDate package (Smith, Brown and Walker, 2018) to extract the 10 most clock-like UCEs. We set up the HKY substitution model for the dataset. We ran two Markov Chain Monte-Carlo (MCMC) with correlated clock rates (Thorne and Kishino, 2002) and 2,000 (burn-in) + 20 (sample frequency) x 20,000 (number of samples) for a total of 402,000 iterations. The convergence of the MCMC has been verified with Tracer v1.7.1 (Rambaut *et al*., 2018) checking for the values of effective sample sizes (ESS), considering ESS > 200 as good convergence for a given parameter. The dated tree has then been used for *Messor* biogeographic inference with DECX, but only one specimen of each taxon was kept.

### 5. Inferring ancestral geographic ranges

To precisely infer the biogeographic origin and diversification of *Messor*, we divided the Palearctic and Afrotropics into 14 areas as follows: 1 - Southern Africa; 2 - Sahelian and Eastern Africa; 3 - Northeast Africa; 4 - Northwest Africa; 5 - Southwest Europe; 6 - Northwest Europe; 7 - Italian Peninsula; 8 - Northeast Europe; 9 - Balkans; 10 - Türkiye and Caucasus; 11 - Arabian Peninsula; 12 - Irano-Indian Asia; 13 - Central Asia; and 14 - Eastern Asia (Fig. S2). We also defined three time periods of 10 million years between the present and 30 Mya.

For the inference, we used DECX (Beeravolu and Condamine, 2016), an extension of the Dispersal-Extinction-Cladogenesis model that can easily handle computational constraints with such many defined areas. The distribution matrix of the investigated *Messor* species (Table S2) has been mainly constructed from the antmaps.org repartition data (Guenard *et al*., 2017), but only when considered as native in a country to discard the introduced species before the historical biogeography inference. The following minor modifications or completions have been made to better suit actual knowledge. The repartition of *M. asmaae*, which is still regarded as an *Aphaenogaster* on antmaps.org despite its recent new combination (Schifani *et al*., 2022), has been completed with the origin of our sample, making it the first mention for Pakistan, and the first mention outside the Arabian Peninsula (NHMUK010363418, BMNH). *Messor meridionalis* was not considered to occur in the Italian peninsula and Southwest Europe because there is a lack of evidence for the presence of this dubious species in these well-studied areas. We assigned *M. bouvieri* to Southwest Europe only, as the species is present in France but only along the Mediterranean coasts (Lebas *et al*., 2016). We discarded mention of *M. sanctus* from Palestine and Syria (Bernard, 1967), as the species has not been mentioned in recent checklists (Vonshak and Ionescu-Hirsch, 2010 for Israel and Tohmé and Tohmé, 1981 for Syria). We discarded *M. aegyptiacus* from the Sahelian and Eastern Africa area as there is only one mention of it in Mauritania (Cagniant and Espadaler, 1998, found in Zouérat Cagniant com. pers.). We assigned the repartition of *M. minor maurus* according to Santschi (1927b), as this subspecies is absent from antmaps.org. The adjacency matrix of the areas has been binarily coded, with 0 whenever two areas are not directly connected at the time considered, and 1 when areas are geographically linked (Table S3).

### 6. Estimating diversification rates

We investigated the diversification rates of *Messor* (58 taxa considered). We first tested whether rates of diversification (speciation minus extinction) remained constant or varied through time, and then tested the impact of palaeoenvironmental variables on diversification rates, using a birth-death likelihood method in which rates may change continuously with an environmental variable, itself varying through time (Condamine, Rolland and Morlon, 2013, 2019). As implemented in RPANDA 2.1 (Morlon *et al*., 2016), we tested four models: (1) a constant-rate birth-death model in which diversification is not associated to the environmental variable; (2) speciation rate varies according to environment and extinction rate does not vary; (3) speciation rate does not vary and extinction rate varies according to environment; and (4) both speciation and extinction rates vary according to environment. We also tested the corresponding models in which speciation and/or extinction are allowed to vary with time, but independently from the environmental variable (time-dependent birth-death models, Morlon, Parsons and Plotkin, 2011). When rates varied with the environment (*E*), we assumed exponential variation, such that λ(E)=λ_0_ × e^αE^ and μ(E)=μ_0_× e^βE^, in which λ_0_ and μ_0_ are the speciation and extinction rates for a given environmental variable for which the value is 0, and α and β are the rates of change according to the environment. Positive values for α or β mean a positive effect of the environment on speciation or extinction (and conversely). As environmental variables, we used palaeotemperature (data retrieved from Westerhold *et al*., 2020, converted to sea-surface temperature by Boschman and Condamine, 2022) and proportion of C_4_ grasses, a proxy for grassland expansion (data retrieved from Kergoat *et al*., 2018). The R package pspline was used to build environmental vectors from the data as input for the birth-death models. In total, we fitted 14 different models to the dated phylogeny of *Messor*. For the list of the tested models and their results, see Table S5. The best-fitting model was selected based on corrected Akaike Information Criterion (AICc) scores and Akaike weight (AICω).

## III. Results

### 1. Phylogeny of the genus *Messor* within the tribe Stenammini

In general, ultrafast bootstrap values of the super-matrix (4,255,038 bp) inferred phylogeny suggest strong support for the main subdivisions of *Stenammini*. Most genera are monophyletic with maximal bootstrap support (Fig. 1). As already inferred in previous work (Branstetter *et al*., 2022), *Goniomma* and *Oxyopomyrmex* were not clearly separated while *Aphaenogaster* species are divided into a tropical clade (named *Deromyrma*, see Branstetter *et al*., 2022) and a Holarctic one (simply named *Aphaenogaster*). In our analysis, *Aphaenogaster* is recovered monophyletic and is the closest relative of *Messor*, a result that contrasts with Branstetter *et al*. (2022) in which *Messor* was nested within *Aphaenogaster*. The monophyly of both genera is supported by maximal bootstrap values. However, the super-tree analysis retrieves *Aphaenogaster* as paraphyletic with *A. subterranea* being the closest relative of *Messor*. gCF confirms phylogenetic conflict among genes, with only 4% of UCEs supporting *Aphaenogaster* as monophyletic. Nevertheless, those 4% of loci appear to carry most of the phylogenetic signal of the dataset, as more than half of the global phylogenetic signal supports *Aphaenogaster* monophyly instead of alternative topologies (sCF of 51.1%).

**Figure 1:**
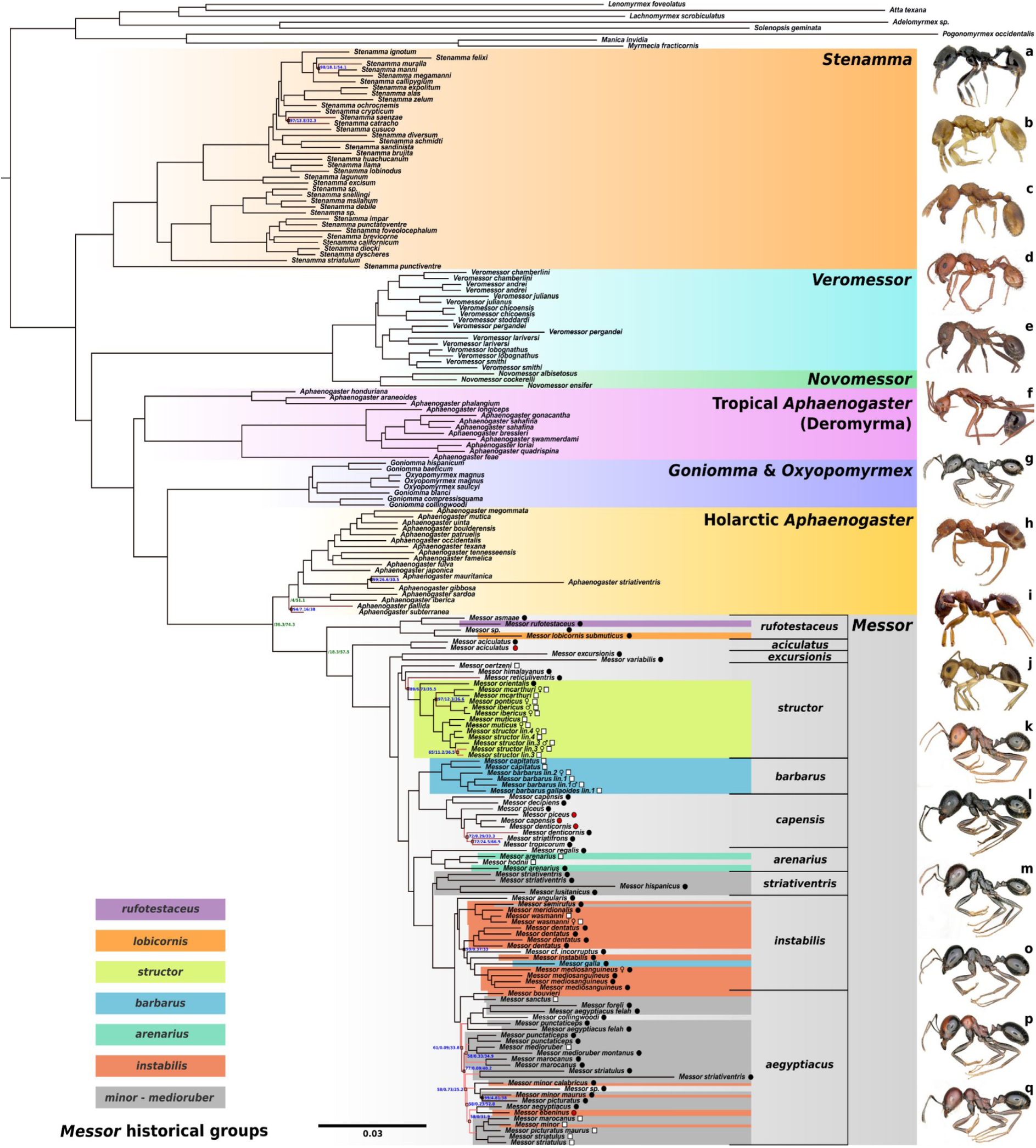
Maximum likelihood phylogeny of the Stenammini tribe. Topology obtained with IQ-TREE on a supermatrix of 2,524 UCEs. Only ultrafast Bootstrap values inferior to 100 are written in blue at nodes, followed by gene (gCF) and site (sCF) concordance factors. Concordance factors of important nodes between Holarctic *Aphaenogaster* and *Messor* are written in green. For the *Messor* genus, circles mean that the sample has been sequenced after capture of UCEs (black circle for new data, red circle for Branstetter et *al.* 2022 data), and white squares mean that the whole genome of the sample has been sequenced. Images of worker specimens (with a code if from antweb.org): a - *Stenamma callipygium* (CASENT0606207, credit to M. Branstetter); b - *S. lagunum* (CASENT0622371, holotype, credit to M. Branstetter); c - *S. californicum* (CASENT0221920, credit to X. Yang); d - *Veromessor lobognathus* (CASENT0104781, credit to A. Nobile); e - *Novomessor albisetosus* (CASENT0102824, credit to A. Nobile); f - *Aphaenogaster swammerdami* (CASENT0421554, credit to A. Nobile); g - *Goniomma blanci* (personal coll., credit to L. Soldati - CBGP); h - *A. uinta* (FMNHINS0000062702, paratype, credit to G. Brilmyer); i - *A. subterranea* (ANTWEB1041832, credit to E. Collins-Sussman); j - *Messor asmaae* (CASENT0922290, credit to M. Esposito); k - *M. oertzeni* (personal coll., credit to L. Soldati - CBGP); l - *M. ibericus* (personal coll., credit to L. Soldati - CBGP); m - *M. barbarus* (personal coll., credit to L. Soldati - CBGP); o - *M. wasmanni* (personal coll., credit to L. Soldati - CBGP); p - *M. picturatus* (personal coll., credit to L. Soldati - CBGP); q - *M. aegyptiacus* (personal coll., credit to L. Soldati - CBGP).

Within *Messor*, the historical *structor* species group (hereafter group) appears monophyletic and supported on both super-matrix and super-tree phylogenies (Fig. 1, Fig S1). However, we included three species (*M. oertzeni*, *M. himalayanus,* and *M. reticuliventris*), which to our knowledge were never mentioned as belonging to this group in recent work (Steiner *et al*., 2018; Barech *et al*., 2020; Salata, Demetriou, *et al*., 2023). In addition to their phylogenetic placement, these three species share the general morphology of the *structor* group (dense pilosity, marked sculpture on head and mesosoma, lack of psammophore, see Bernard, 1955). Other congruently supported clades are observed, but without having been specifically defined until now. Based on phylogenetic evidence, we propose to name *capensis* group for the Southern Afrotropical *Messor* clade, and *excursionis* group for *M. excursionis* and *M. variabilis* clade. The *aciculatus* group is only composed of one species considering our sampling, as its phylogenetic position is clearly apart from the rest of the tree. We extend the *rufotestaceus* group with *M. asmaae*. This result confirms the recent morphological revision and genus reassignment of *Aphaenogaster asmaae* Sharaf, 2018 into *Messor* (Schifani *et al*., 2022). *Messor lobicornis submuticus* also appears in our extended *rufotestaceus* group despite being morphologically divergent. The same occurs in the *arenarius* group with *M. hodnii*, which is surprisingly nested between two *M. arenarius* specimens. We also included *M. regalis* in the group to form a slightly anterior clade. In our sampling, the historical *barbarus* group was only represented by three species, but *M. galla* does not group with the two others (*M. barbarus* and *M. capitatus*) and is thus excluded from the *barbarus* group.

Groups *minor-medioruber* and *instabilis*, used in Barech *et al*. (2020) and Salata, Lapeva-Gjonova, *et al*. (2023) respectively, do not constitute clades. This result was expected as these two groups have overlapping species composition in the aforementioned references. The *minor*-*medioruber* group is divided into a clade containing species from the Iberian peninsula and Northwest Africa (*M. hispanicus*, *M. lusitanicus* and *M. striativentris*); and a large clade of mostly North-West African species. We renamed these two clades the *striativentris* and *aegyptiacus* group, respectively. More particularly, the *aegyptiacus* group is a large clade of closely related species/subspecies with diverse morphologies and poorly supported relationships. Further work might help to subdivide it into several species complexes. The historical *instabilis* group is one of the most anciently used and species-rich groups if considered to its greater extent (Santschi, 1927a). It appears that the group supported by our phylogeny contains almost all the species considered by Salata, Lapeva-Gjonova, *et al*. (2023) for which we had material, with the exception of *M. bouvieri*, *M. ebeninus,* and *M. minor* + *ssp.* that instead belong to the *aegyptiacus* group.

In our phylogeny, the following species appear to be non-monophyletic: *M. capensis*, *M. piceus*, *M. denticornis, M. striatulus*, *M. strativentris*, *M. punctaticeps*, *M. arenarius*, *M. aegyptiacus felah*, *M. marocanus*, *M. minor* + *ssp*. and *M. picturatus* + *ssp*. While this result might be due to technical artifacts or identification inconsistencies between taxonomists, we tested whether it might be due to the presence of hybrid individuals in our dataset.

### 2. F1 Hybrid detection within the tribe Stenammini

To check whether some of the sequenced individuals are hybrids, we computed the γ/θ ratio of each individual following an approach designed to detect F1 hybrids from single individual genetic data (Weyna, Romiguier and Mullon, 2021). Individuals with γ/θ ratio superior to 0.557 (*i.e.* ratio value of a *Paratrechina longicornis* worker known as an F1 hybrid, Weyna, Romiguier and Mullon, 2021) were considered as potential hybrids. Results on our dataset retrieved 9 samples as hybrid candidates (see Fig. 2), including 6 from UCE-capture samples and 3 from WGS samples. These 9 samples include *M. barbarus*, which is known for practicing social hybridogenesis, a reproductive system involving workers that are exclusively hybrids (Romiguier *et al*., 2017). Interestingly, this also includes a worker from the subspecies *M. barbarus gallaoides*, a morphological variant from North Africa that also appears as unambiguously hybrid. Three other *Messor* workers also appear as candidates. Among them, one individual of *M. denticornis* and *M. capensis* are retrieved. This could match with the inconsistencies displayed in the phylogeny, where the two hybrid individuals do not respectively group with their non-hybrid conspecifics. As their supposed hybridism may be responsible for this incongruent clustering, we decided to remove these individuals from the dating and diversification analysis. The other hybrid retrieved among the genus is *M. hodnii*, which has been inferred from WGS data. Its ratio mean is just above the ratio threshold, and once again this could explain why our sequenced specimen appears misplaced, nested within *M. arenarius* individuals. Outside of these 5 *Messor* candidates, the other hybrids retrieved have been found in the genera *Veromessor*, *Goniomma*, *Aphaenogaster*, and the clade *Deromyrma*. *Veromessor julianus* is the only one represented by more than one specimen, however, they do cluster together in the phylogeny.

**Figure 2:**
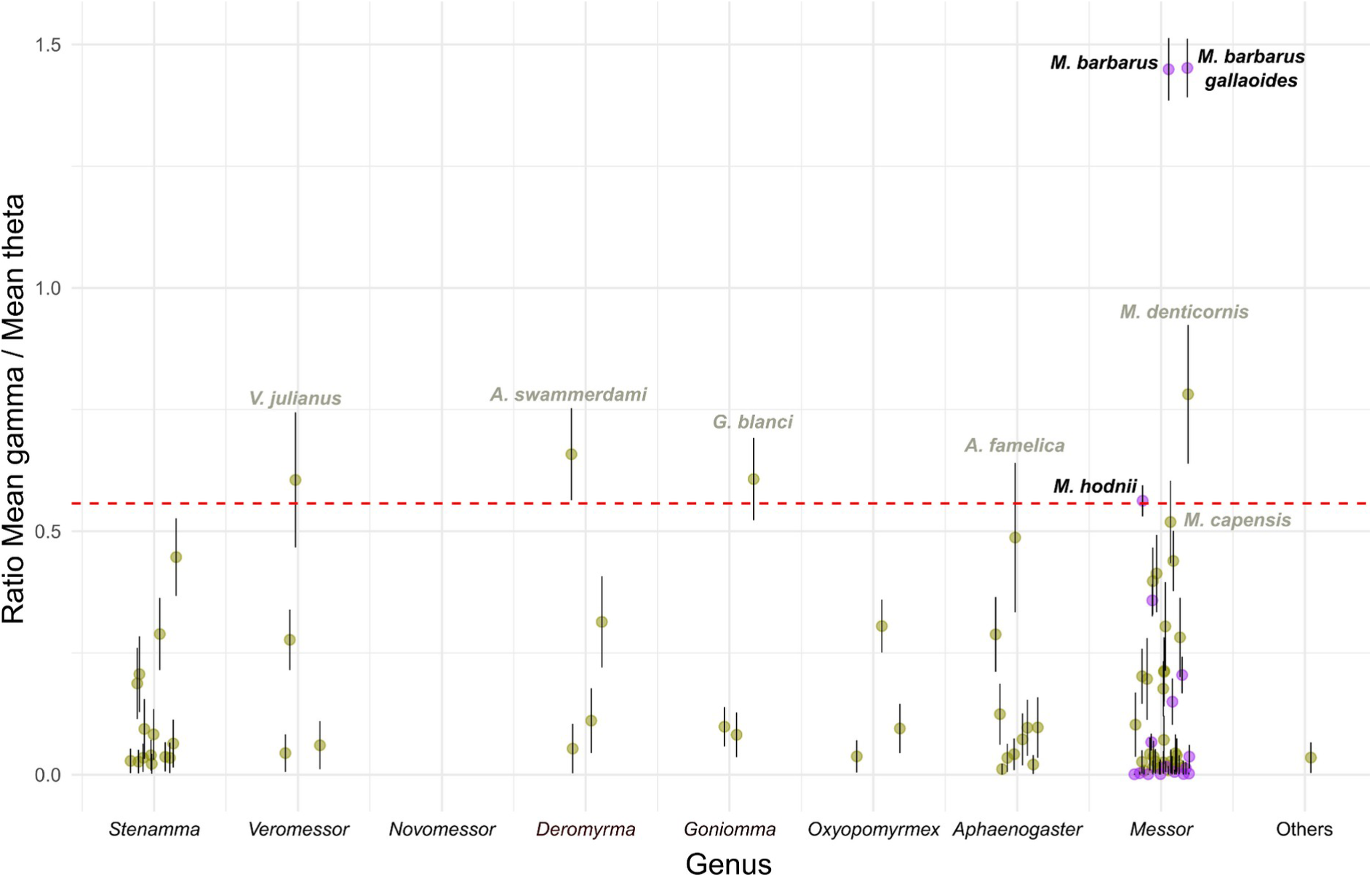
Heterozygosity scan of investigated specimens with a contamination filter. The estimate is the ratio between the divergence parameter γ and the ancestral population mutation rate θ. Purple points are ratios estimated with BUSCO genes on WGS specimens, whereas yellow points come from UCE capture. The red dashed line estimates a threshold for abnormal heterozygosity, as it passes by our lowest ratio value of already sequenced hybridogenetic species, *Paratrechina longicornis*. Points above the threshold are candidates for hybridism. Only the labels of points above the line, or with error bars crossing it, are displayed. Points with mean theta values inferior to 0.0005 have been discarded from the plot.

### 3. Age and historical biogeography of *Messor*

The Bayesian inferences of divergence times with MCMCtree converged well with most parameters having ESS values above 200. The divergence time between the genus *Aphaenogaster* and the genus *Messor* was estimated at 21.68 Mya (95% HPD: 22.97 - 20.39 Mya). The age of the diversification of *Messor* is inferred at 19.67 Mya (95% HPD: 21.13 - 18.22 Mya; Fig. 3).

**Figure 3:**
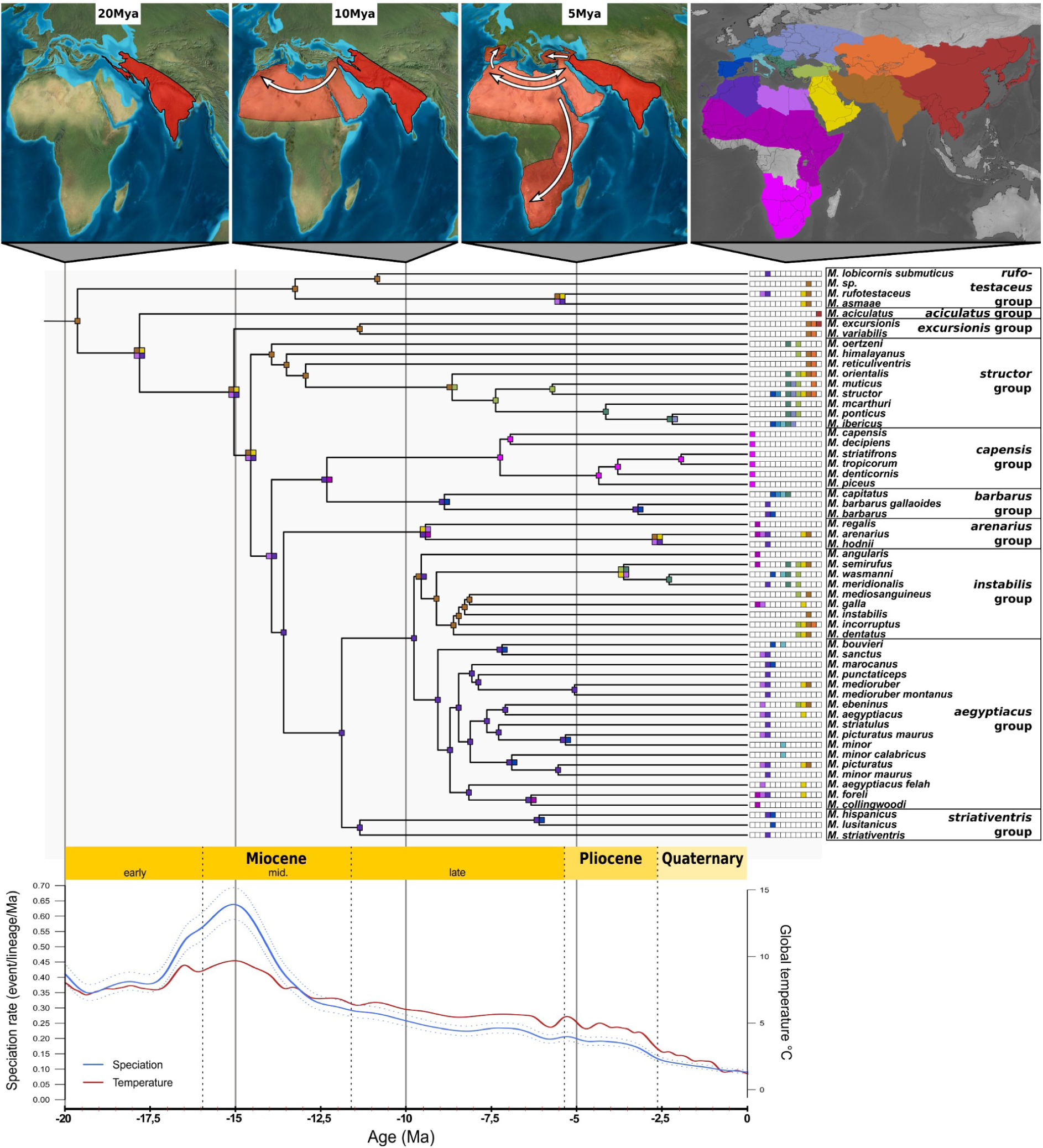
Dendrogram of historical biogeography of the *Messor* genus and its taxonomic groups with temperature-correlated diversification rates and the illustration of major dispersal routes. Selection of only one representative for each taxon has been made depending on the best loci-covered individual.

The biogeographic analyses estimated an Irano-Indian range as the ancestral area for *Messor* (Fig. 3). Our results suggested a rapid geographic dispersion from their origin area towards the Western Palearctic, reaching Northeastern Africa during the early Miocene. We inferred a major diversification event, during the middle Miocene climatic optimum (MMCO: 17-14 Mya; Steinthorsdottir *et al*., 2021). During this climatic event, we inferred the emergence of the common ancestors of all *Messor* groups except for the *rufotestaceus* and *aciculatus* groups (which diverged earlier), around the latitudinal axis from Northwest Africa to Irano-Indian region. After the middle Miocene climatic transition (MMCT; 14 Mya), where temperatures started to decrease, the *arenarius* group emerged from Northwest Africa. The end of the middle Miocene saw then the emergence of *capensis* and *barbarus* groups from the same ancestral clade, as well as the *striativentris* group, in the northern half of Africa.

Finally, the two last groups, *instabilis* and *aegyptiacus*, differentiated during the middle Miocene from a common ancestor in Northwest Africa, approximately 12 Mya (Fig. 3). We found that Europe has been reached later, a few million years later in the late Miocene, from Northeast Africa to Southwest Europe with the origin of the *barbarus*, *aegyptiacus* and *striativentris* groups; and later during the Pliocene, from Türkiye to the Balkans with the diversification of *structor* and *instabilis* groups.

### 4. Diversification pattern of *Messor*

Fitting a series of birth-death models to the phylogeny of *Messor* indicated that the null hypothesis of constant diversification rates can be rejected, suggesting that rates have varied through time (Table S5). The rate variation was better explained by temperature fluctuations rather than time with a model assuming temperature-dependent speciation and no extinction receiving the lowest AICc and highest AICω (AICω = 0.522, ΔAICc > 2 with the second best model; Table S5). Maximum-likelihood parameter estimates of the best model indicated a positive correlation (α = 0.3) with speciation rate for the clade, suggesting higher speciation rates during warm periods. This translates into faster rates of speciation during the middle Miocene for instance, and lower speciation rates in the Pleistocene (Fig. 3). Note that the second best model was a model with temperature-dependent speciation and constant extinction estimated to be close to zero.

## IV. Discussion

### 1. Phylogenomics

Previous work retrieved the *Messor* genus as potentially nested within a Holarctic *Aphaenogaster* clade (hereafter, abbreviated as *Aphaenogaster* in contrast to the *Aphaenogaster* tropical clade renamed *Deromyrma*, Branstetter *et al*., 2022). Here, our phylogeny separates *Messor* and *Aphaenogaster* genera with 100% ultrafast bootstrap support. This difference likely stems from the high number of *Messor* taxa in our dataset, an important improvement compared to previous work (6 vs 51 species here), representing approximately a third of the currently known diversity. However, a major limitation of our dataset is the low number of *Aphaenogaster* species, especially in groups originating from the Palearctic bioregion. Further studies with a higher number of *Aphaenogaster* species will be necessary to confirm that both genera are consistently separated.

Greater taxa depth and redundancy could have been useful to clarify poorly supported relationships or to better recognize potential wrong identifications causing inconsistent species clustering. Nevertheless, the main species groups are not expected to change drastically, as most historical taxonomic groups are already represented. The majority of ambiguous relationships within *Messor* stem in the *instabilis* and *aegyptiacus* groups, whose origin seems to be in North-West Africa. This is not surprising as it is a major diversity hotspot of the genus. This species-rich region might have promoted rapid and numerous speciation events, leading to short branches difficult to resolve through phylogenetic inference. Resolving these relationships might be additionally hampered by the use of UCE loci, as these conserved regions may lack enough differences for separating closely related species that diversified after fast and successive speciation events.

Contrary to expectations, some individuals thought to belong to the same species do not always cluster together. This could be due to wrong identifications, which are not uncommon for this genus, or due to few informative substitutions, as suggested by low node support values. However, some cases are interesting, like *M. hodnii* which renders *M. arenarius* paraphyletic. Those two species cannot be morphologically confused as they are strongly unalike. Little is known about *M. hodnii*, as its discovery has been made recently (Barech *et al*., 2020). Another interesting case is observed with incongruent taxa clustering for Southern Africa *Messor* species. This specifically includes *M. capensis*, *M. decipiens* and *M. piceus*, which do not seem to be monophyletic. Although this could be explained by wrong identifications, it was mentioned before that these extremely similar species could in fact be the same species (Bolton, 1982). If this is indeed the case, *M. piceus* Stitz, 1923 and *M. decipiens* Santschi, 1917 need to be synonymized with *M. capensis* Mayr, 1862. *M. denticornis* also appears polyphyletic, however in this case, incorrect identification would be surprising, as this species should be reliably distinguishable from its relatives through eye size (Bolton, 1982). The most likely explanation for such inconsistencies is a methodological artifact due to heterogeneous data origin. Indeed, we note that *Messor* individuals with genetic data retrieved from the literature (Branstetter *et al*., 2022) cluster together, in a separate group than individuals of the same species with genetic data produced in this study. This could be explained because the later study has targeted the capture of 1,510 UCE loci whereas we have targeted 2,524 UCE loci, which is likely to imply a non-random distribution of missing data and artifactual clustering among closely related specimens.

We suggest that the investigated *M. meridionalis* specimen, because of its morphological inspection, its dubious status (*incertae sedis* taxon, Borowiec, 2014; Salata and Borowiec, 2019; Kiran and Karaman, 2021; Salata, Lapeva-Gjonova, *et al*., 2023), its sampling origin (Yougoslavia, 1986, NHMUK010363380, BMNH) and its phylogenetic position, should correspond to *M. wasmanni*.

*Messor lobicornis submuticus* appears in our extended *rufotestaceus* group, but its morphological divergence and low UCE coverage (118/2,524) make this result uncertain. Moreover, the existence and historical use of the morphological *lobicornis* group would need to be genetically investigated, with a better sampling to infer its systematics properly.

Regarding unidentified *Messor* species, the *M. sp.* clustering in the *aegyptiacus* group could not be properly identified due to the small size of the specimen. However, the one clustering in the *rufotestaceus* group, which comes from a pin series of three workers sampled from Iraq (NMUK010363417, BMNH), corresponds the most to *M. thoracica* (Mayr, 1862), a junior synonym of *M. rufotestaceus*. We believe this taxon should be restored under the correct spelling *M. thoracicus* as its synonymization seems arbitrary, but further work will be needed to prove it.

Hybrid scan results confirm the social hybridogenesis of *Messor barbarus* (Romiguier *et al*., 2017) in extra-European samples (investigated specimens come from Morocco and Algeria), as for the subspecies *M. barbarus gallaoides,* which is newly reported as hybrid. Whereas the taxa hybridizes with another subspecific lineage or with one of the already known *M. barbarus* lineage is still to be known. Among other WGS samples, only *M. hodnii* stands out as a potential hybrid. For UCE-capture samples, 6 specimens could be or are above the threshold, however, these results should be interpreted with caution as UCE data may lack enough loci and polymorphism to reliably infer hybrid status, whereas BUSCO has been shown to be more efficient for heterozygosity assessment (Weyna *et al*., 2022). The 6 UCE capture specimens feature only from 796 to 875 UCE loci, which represent a maximum of 956,114 bp, whereas the 3 WGS specimens have 1925 to 1948 UCE loci, and 4027 to 4156 BUSCO loci (Table S1). This calls for further studies in these species to explore a potentially unusual reproductive system such as social hybridogenesis. It is to note that the sequenced *M. ebeninus* does not appear as hybrid, as it would be expected (Romiguier *et al*., 2017). This might be due to a missidentification, or if the specimen was an alate, which are of pure lineage because of their social hybridogenesis.

### 2. Origin, biogeographic history and diversification

The inferred ancestral area of the *Messor* genus corresponds to Irano-Indian Asia in the early Miocene, approximately 20 Mya. This origin suggests that the evolution of this genus might be linked to climatic and environmental changes induced by the Himalayan orogenic process that was still ongoing at this time (Ji *et al*., 2020) and may have contributed to the isolation and emergence of the genus (see Tardif *et al*., 2023). Moreover, this major geological event is suspected to have promoted C4 grasses spread due to the progressive aridification of the region (Wu *et al*., 2014), which is an important source of food for *Messor* ants due to their granivorous diet.

Regarding our results, the *Messor* genus likely reached North Africa during its early history. This first and rapid dispersal route out of their ancestral area could help understand the current observed diversity hotspots. Indeed, North-West Africa and especially the Middle-East are geographic crosspoints that could have allowed repeated migrations, which likely contributed to the high diversity of these regions. Furthermore, the arid climate and mountainous topography—such as the Atlas range in North-West Africa and the Zagros, Alborz, Taurus, and Pontic ranges in the Middle-East—may have created ideal conditions for speciation and diversification of *Messor* species.

Regarding Europe, two potential reasons may explain why *Messor* species arrived in the region later than in others. The first one is the connectivity of the continent, which was not directly reachable from the time and emergence area of the *Messor* genus (see Table S2). The second one is related to its latitude with a colder climate, thus limiting the grassland dominance to Mediterranean coasts. However, two different dispersal routes can be reported towards the continent. The first to have been achieved approximately 9 Mya comes from the Gibraltar strait, and involves the *barbarus* group. The other route from Türkiye towards the Balkans occurred later, nearly 4 Mya, and implies most of the *structor* group. Glaciation events and the lowering of global sea levels due to ice cap formation have likely influenced European migrations. Because of the geographic bottleneck of the Gibraltar strait between Africa and Europe, and of the Bosphorus and Dardanelles straits between Asia and Europe, crossings might have been temporarily impeded or allowed depending on the sea level and land connectivity, thus contributing to successive vicariance and speciation events (Mas-Peinado *et al*., 2022). Examples of intermittent Gibraltar-crossing dynamics, as hypothesized for other ants (Tinaut and Ruano, 2021), could coincide with several *Messor* species pairs: (i) *M. hispanicus* / *M. lusitanicus*, which are closely related and on both sides of Gibraltar strait (*M. lusitanicus* is considered as exotic in Morocco, based on antmaps.org); (ii) *M. bouvieri* / *M. sanctus*, which are also sister species, but the first one is only present in Europe while the second is only present in North-Africa. Whereas the latter example could illustrate a typical allopatric speciation, the observed sympatry of *M. hispanicus* and *M. lusitanicus* cannot exclude other speciation types in which the Gibraltar strait may not be involved.

Other minor dispersal routes can be noticed towards Southern Africa and Central Asia. The first one is clearly depicted as it has led to the *capensis* group, probably passing through the East African “arid corridor” during the late Miocene (Bobe, 2006). This matches with an untested hypothesis that *M. regalis* and *M. cephalotes*, two Afrotropical species, came from the Nile Valley due to their relatedness with *M. arenarius* (Santschi, 1938). The second minor route is not that obvious due to our incomplete diversity sampling for these areas but is expected to have lead to most if not all of the Central and East Asian diversity, circling the Himalayan range from the West towards Northern steppes and Gobi desert, where we suspect an undescribed diversity could still await description.

These results should nevertheless be taken with caution. A possible bias of our analysis is potential species misidentifications when selecting one representative individual per species/subspecies. This could lead to wrong inferred areas, as well as poor taxonomic grouping representation. Moreover, we still lack taxa depth to better define biogeographic history dynamics and their taxonomic consequences. Our sampling is biased towards the occidental diversity despite the genus originating in Asia. Also, we still lack crucial knowledge about some species delimitations and relevance, especially since hybridization can occur during obligatory or facultative interspecific interactions (Steiner *et al*., 2011; Romiguier *et al*., 2017). Another linked problem is the reliability of AntMaps (Guenard *et al*., 2017) repartition data for area assignment, as in the cryptic species complex unraveled from *M. structor* (Steiner *et al*., 2018). The latter species has many old and unrevised mentions that may often designate *M. ponticus*, *M. muticus*, *M. mcarthuri* or *M. ibericus*. However, all these issues are limited to recent species history details, but are unlikely to have affected the overall biogeographic history of the *Messor* genus.

Interestingly, this history appears consistent with a specialized ecology into arid environments, as the initial dispersal route of the genus appears to have followed progressive aridification around the Indo-Iranian region (Fig. 3). This pattern is strikingly similar to what has been recently inferred for the *Cataglyphis* genus, which is considered the other major genus of desertic ants in the Palearctic. Both genera emerged approximately 20 Mya between the Middle-East and South Asia then dispersed towards Europe and Northern Africa (Lecocq De Pletincx *et al*., 2024). Remarkably, the parallel biogeographical histories of *Messor*, reconstructed here from molecular data, and *Cataglyphis*, were predicted over a century ago (Emery, 1912, 1920; Kugler, 1988). The similar dispersal routes of these two desertic ant genera reinforce the idea that species specialized for extreme environments have evolved more stereotypically in response to past climate changes compared to generalist species. Such trends may extend to their responses to global warming, making predictions for extreme specialists more straightforward. Further research evaluating how past and ongoing global warming affects these biome specialists may enhance our understanding of how climate change differentially impacts species.

## Supplementary Material

**Figure S1.**
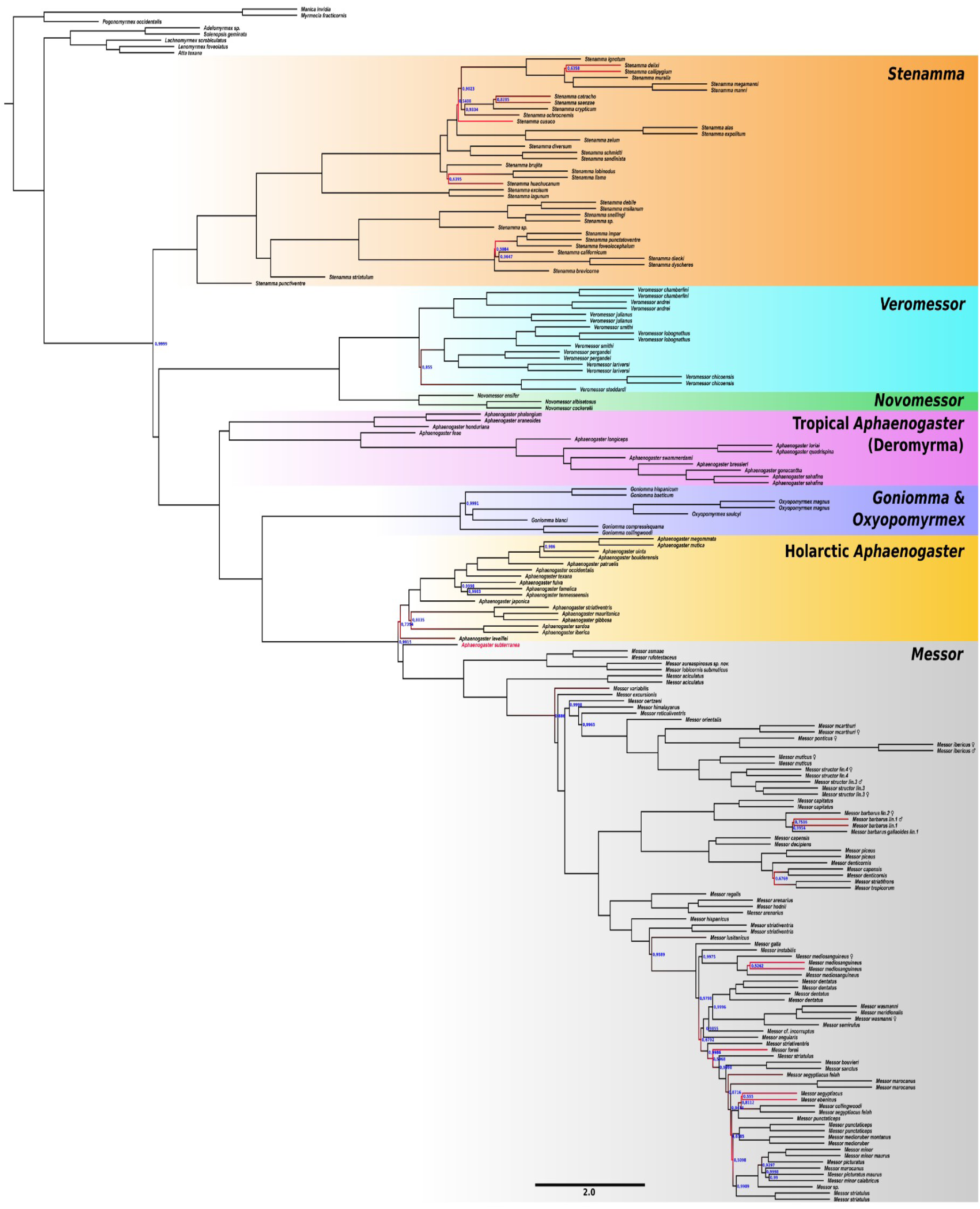
Consensus tree of 2203 gene (UCEs) trees inferred with wASTRAL-hybrid.

**Figure S2.**
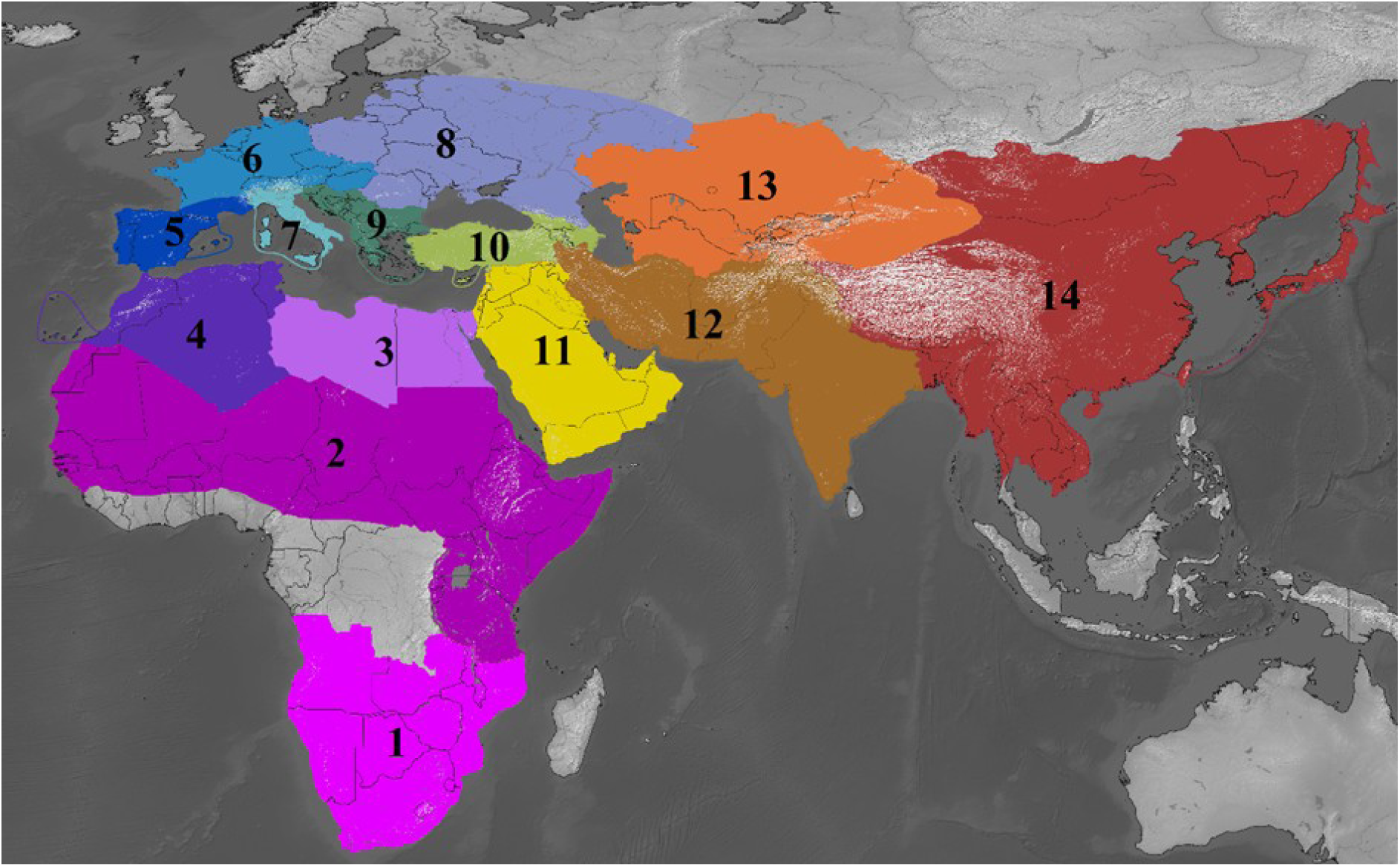
Map of the areas defined for the historical biogeography. 1 - Southern Africa; 2 - Sahelian and Eastern Africa; 3 - Northeast Africa; 4 - Northwest Africa; 5 - Southwest Europe; 6 - Northwest Europe; 7 - Italian Peninsula; 8 - Northeast Europe; 9 - Balkans; 10 - Türkiye and Caucasus; 11 - Arabian Peninsula; 12 - Irano-Indian Asia; 13 - Central Asia; 14 - Eastern Asia. Mountain ranges, which can be major dispersal barriers, can be perceived inside the defined zones with whiter tones.

**Table S1.**
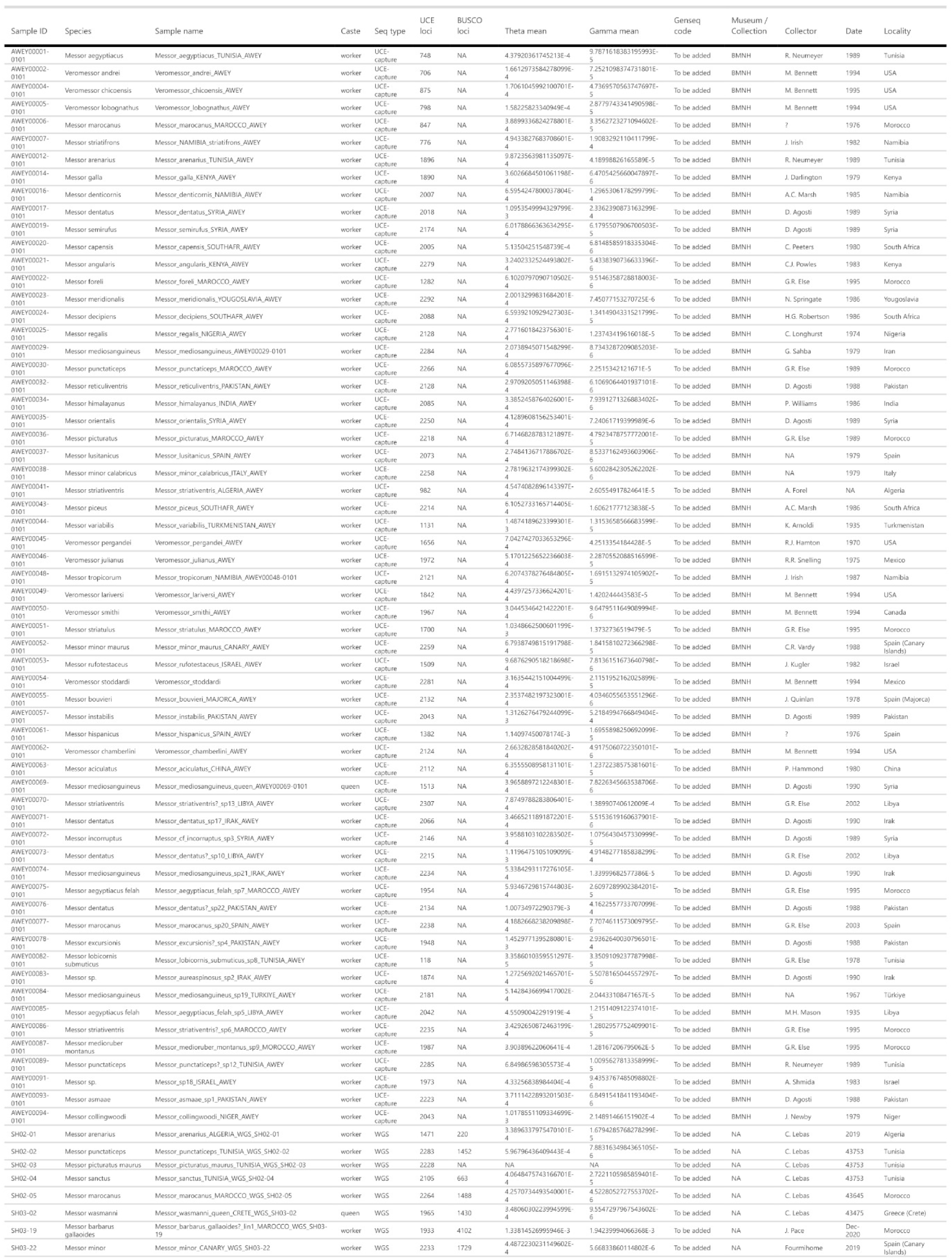
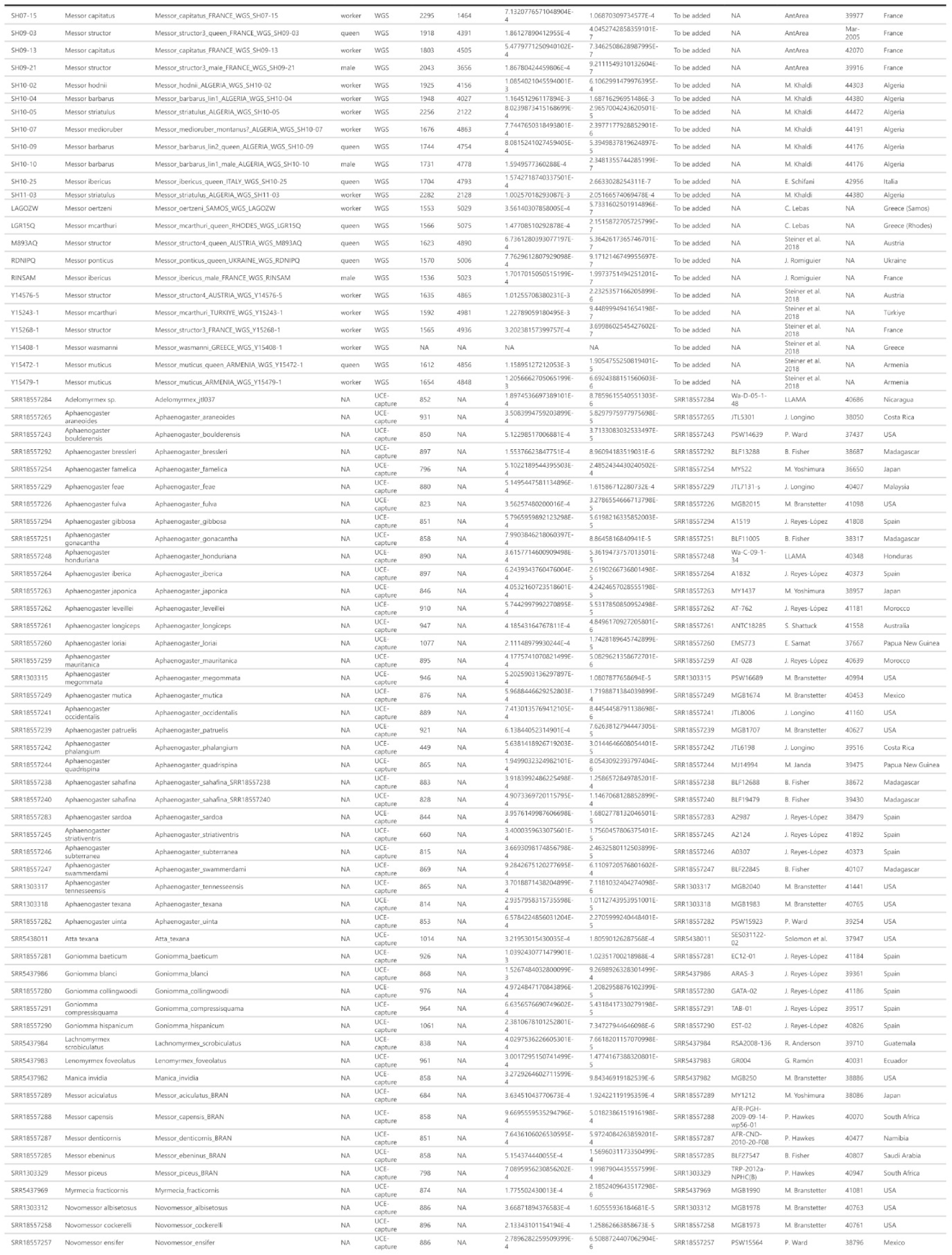
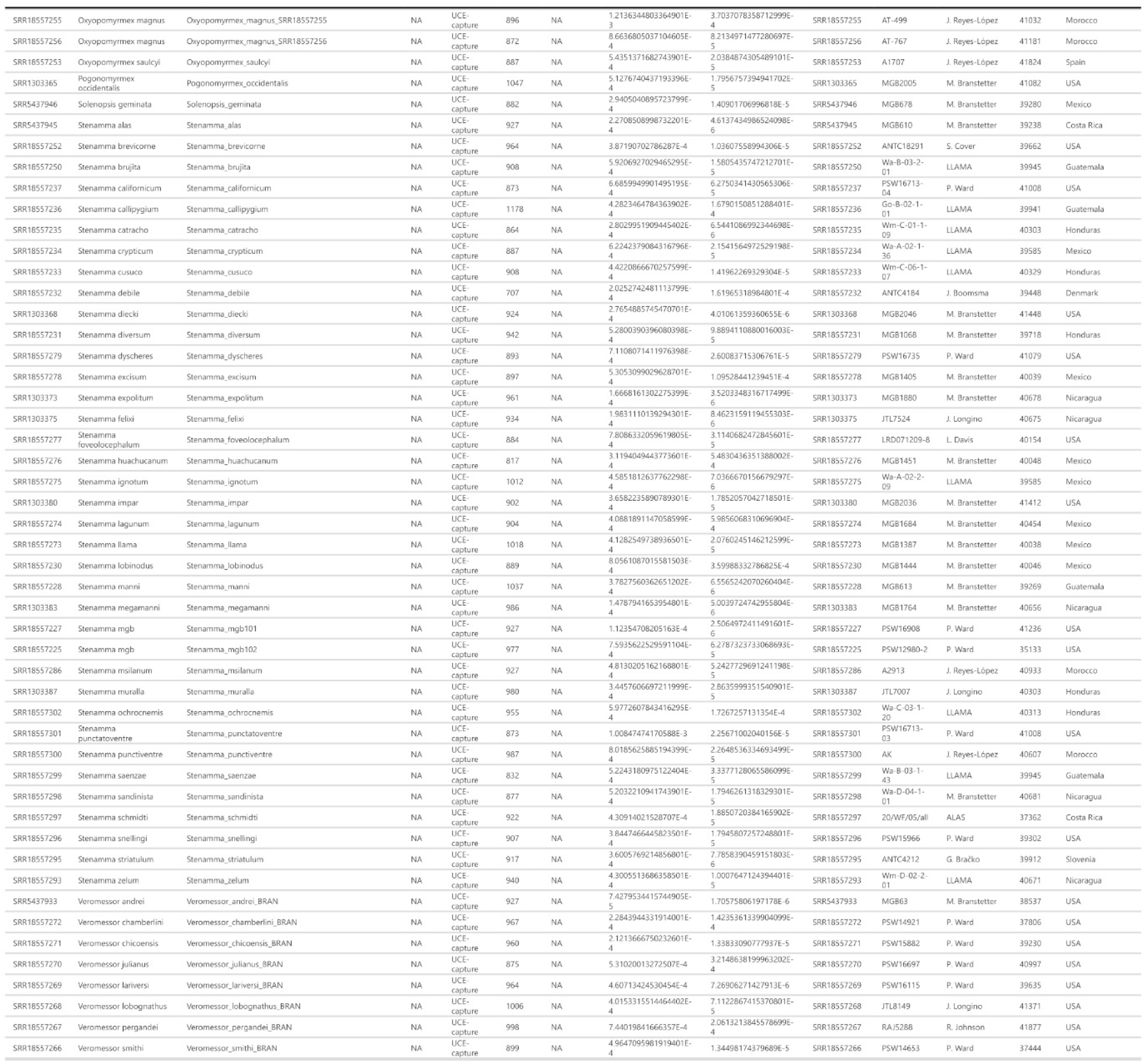
Information for the taxon sampling of the study. The table shows for each sample, its species attribution, the number of sequenced loci, the estimated parameters for the heterozygosity scan (theta and gamma), the voucher code from the Museum, and the locality and data of collection.

**Table S2.** Distribution matrix of *Messor* species among the defined biogeographic areas.

| *Messor* Species | 1 -<br>Southern<br>Africa | 2 -<br>Sahelian<br>and<br>Eastern<br>Africa | 3 -<br>North<br>East<br>Africa | 4 -<br>North<br>West<br>Africa | 5 -<br>South<br>West<br>Europe | 6 -<br>North<br>West<br>Europe | 7 -<br>Italian<br>Peninsula | 8 -<br>North<br>East<br>Europe | 9 -<br>Balkans | 10 -<br>Türkiye<br>and<br>Caucasus | 11 -<br>Arabian<br>Peninsula | 12 -<br>Irano-Indian<br>Asia | 13 -<br>Central<br>Asia | 14 -<br>Eastern<br>Asia |
| --- | --- | --- | --- | --- | --- | --- | --- | --- | --- | --- | --- | --- | --- | --- |
| *aciculatus* | 0 | 0 | 0 | 0 | 0 | 0 | 0 | 0 | 0 | 0 | 0 | 0 | 0 | 1 |
| *aegyptiacus* | 0 | 0 | 1 | 1 | 0 | 0 | 0 | 0 | 0 | 0 | 1 | 0 | 0 | 0 |
| *aegyptiacus felah* | 0 | 0 | 1 | 0 | 0 | 0 | 0 | 0 | 0 | 0 | 1 | 0 | 0 | 0 |
| *angularis* | 0 | 1 | 0 | 0 | 0 | 0 | 0 | 0 | 0 | 0 | 0 | 0 | 0 | 0 |
| *arenarius* | 0 | 1 | 1 | 1 | 0 | 0 | 0 | 0 | 0 | 0 | 1 | 1 | 0 | 0 |
| *asmaae* | 0 | 0 | 0 | 0 | 0 | 0 | 0 | 0 | 0 | 0 | 1 | 1 | 0 | 0 |
| *barbarus* | 0 | 0 | 0 | 1 | 1 | 0 | 0 | 0 | 0 | 0 | 0 | 0 | 0 | 0 |
| *barbarus gallaoides* | 0 | 0 | 0 | 1 | 0 | 0 | 0 | 0 | 0 | 0 | 0 | 0 | 0 | 0 |
| *bouvieri* | 0 | 0 | 0 | 0 | 1 | 1 | 1 | 0 | 0 | 0 | 0 | 0 | 0 | 0 |
| *capensis* | 1 | 0 | 0 | 0 | 0 | 0 | 0 | 0 | 0 | 0 | 0 | 0 | 0 | 0 |
| *capitatus* | 0 | 0 | 0 | 0 | 1 | 1 | 1 | 0 | 1 | 0 | 0 | 0 | 0 | 0 |
| *collingwoodi* | 0 | 1 | 0 | 0 | 0 | 0 | 0 | 0 | 0 | 0 | 0 | 0 | 0 | 0 |
| *decipiens* | 1 | 0 | 0 | 0 | 0 | 0 | 0 | 0 | 0 | 0 | 0 | 0 | 0 | 0 |
| *dentatus* | 0 | 0 | 0 | 0 | 0 | 0 | 0 | 0 | 0 | 1 | 1 | 1 | 0 | 0 |
| *denticornis* | 1 | 0 | 0 | 0 | 0 | 0 | 0 | 0 | 0 | 0 | 0 | 0 | 0 | 0 |
| *ebeninus* | 0 | 0 | 1 | 0 | 0 | 0 | 0 | 0 | 0 | 1 | 1 | 1 | 0 | 0 |
| *excursionis* | 0 | 0 | 0 | 0 | 0 | 0 | 0 | 0 | 0 | 0 | 0 | 1 | 1 | 1 |
| *foreli* | 0 | 1 | 1 | 1 | 0 | 0 | 0 | 0 | 0 | 0 | 1 | 0 | 0 | 0 |
| *galla* | 0 | 1 | 1 | 0 | 0 | 0 | 0 | 0 | 0 | 0 | 1 | 0 | 0 | 0 |
| *himalayanus* | 0 | 0 | 0 | 0 | 0 | 0 | 0 | 0 | 0 | 1 | 0 | 1 | 1 | 0 |
| *hispanicus* | 0 | 0 | 0 | 1 | 1 | 0 | 0 | 0 | 0 | 0 | 0 | 0 | 0 | 0 |
| *hodnii* | 0 | 0 | 0 | 1 | 0 | 0 | 0 | 0 | 0 | 0 | 0 | 0 | 0 | 0 |
| *ibericus* | 0 | 0 | 0 | 0 | 1 | 1 | 1 | 1 | 1 | 0 | 0 | 0 | 0 | 0 |
| *incorruptus* | 0 | 0 | 0 | 0 | 0 | 0 | 0 | 0 | 0 | 1 | 1 | 1 | 1 | 0 |
| *instabilis* | 0 | 0 | 0 | 0 | 0 | 0 | 0 | 0 | 0 | 0 | 0 | 1 | 0 | 0 |
| *lamellicornis* | 0 | 0 | 0 | 0 | 0 | 0 | 0 | 0 | 0 | 0 | 0 | 0 | 1 | 0 |
| *lobicornis submuticus* | 0 | 0 | 0 | 1 | 0 | 0 | 0 | 0 | 0 | 0 | 0 | 0 | 0 | 0 |
| *lusitanicus* | 0 | 0 | 0 | 0 | 1 | 0 | 0 | 0 | 0 | 0 | 0 | 0 | 0 | 0 |
| *marocanus* | 0 | 0 | 0 | 1 | 1 | 0 | 0 | 0 | 0 | 0 | 0 | 0 | 0 | 0 |
| *mcarthuri* | 0 | 0 | 0 | 0 | 0 | 0 | 0 | 0 | 1 | 1 | 0 | 0 | 0 | 0 |
| *medioruber* | 0 | 0 | 1 | 1 | 0 | 0 | 0 | 0 | 0 | 0 | 1 | 1 | 0 | 0 |
| *medioruber montanus* | 0 | 0 | 0 | 1 | 0 | 0 | 0 | 0 | 0 | 0 | 0 | 1 | 0 | 0 |
| *mediosanguineus* | 0 | 0 | 0 | 0 | 0 | 0 | 0 | 0 | 0 | 1 | 0 | 1 | 0 | 0 |
| *meridionalis* | 0 | 0 | 0 | 1 | 0 | 0 | 0 | 0 | 1 | 1 | 0 | 0 | 0 | 0 |
| *minor* | 0 | 0 | 0 | 0 | 0 | 0 | 1 | 0 | 0 | 0 | 0 | 0 | 0 | 0 |
| *minor calabricus* | 0 | 0 | 0 | 0 | 0 | 0 | 1 | 0 | 0 | 0 | 0 | 0 | 0 | 0 |
| *minor maurus* | 0 | 0 | 0 | 1 | 0 | 0 | 0 | 0 | 0 | 0 | 0 | 0 | 0 | 0 |
| *muticus* | 0 | 0 | 0 | 0 | 0 | 0 | 0 | 1 | 1 | 1 | 0 | 0 | 1 | 0 |
| *oertzeni* | 0 | 0 | 0 | 0 | 0 | 0 | 0 | 0 | 1 | 1 | 0 | 0 | 0 | 0 |
| *orientalis* | 0 | 0 | 0 | 0 | 0 | 0 | 0 | 0 | 1 | 1 | 1 | 1 | 1 | 0 |
| *piceus* | 1 | 0 | 0 | 0 | 0 | 0 | 0 | 0 | 0 | 0 | 0 | 0 | 0 | 0 |
| *picturatus* | 0 | 0 | 1 | 1 | 0 | 0 | 0 | 0 | 0 | 0 | 1 | 1 | 0 | 0 |
| *picturatus maurus* | 0 | 0 | 1 | 1 | 0 | 0 | 0 | 0 | 0 | 0 | 0 | 0 | 0 | 0 |
| *ponticus* | 0 | 0 | 0 | 0 | 0 | 0 | 0 | 1 | 1 | 1 | 0 | 0 | 0 | 0 |
| *punctaticeps* | 0 | 0 | 0 | 1 | 0 | 0 | 0 | 0 | 0 | 0 | 0 | 0 | 0 | 0 |
| *regalis* | 0 | 1 | 0 | 0 | 0 | 0 | 0 | 0 | 0 | 0 | 0 | 0 | 0 | 0 |
| *reticuliventris* | 0 | 0 | 0 | 0 | 0 | 0 | 0 | 0 | 0 | 0 | 0 | 1 | 1 | 0 |
| *rufotestaceus* | 0 | 0 | 1 | 1 | 0 | 0 | 0 | 0 | 0 | 0 | 1 | 1 | 0 | 0 |
| *sanctus* | 0 | 0 | 1 | 1 | 0 | 0 | 0 | 0 | 0 | 0 | 1 | 0 | 0 | 0 |
| *semirufus* | 0 | 1 | 0 | 0 | 0 | 0 | 0 | 0 | 1 | 1 | 1 | 1 | 0 | 0 |
| *sp. cf. thoracica* | 0 | 0 | 0 | 0 | 0 | 0 | 0 | 0 | 0 | 0 | 0 | 1 | 0 | 0 |
| *striatifrons* | 1 | 0 | 0 | 0 | 0 | 0 | 0 | 0 | 0 | 0 | 0 | 0 | 0 | 0 |
| *striativentris* | 0 | 0 | 0 | 1 | 0 | 0 | 0 | 0 | 0 | 0 | 0 | 0 | 0 | 0 |
| *striatulus* | 0 | 0 | 0 | 1 | 0 | 0 | 0 | 0 | 0 | 0 | 0 | 0 | 0 | 0 |
| *structor* | 0 | 0 | 0 | 0 | 1 | 1 | 0 | 1 | 1 | 1 | 1 | 1 | 1 | 0 |
| *tropicorum* | 1 | 0 | 0 | 0 | 0 | 0 | 0 | 0 | 0 | 0 | 0 | 0 | 0 | 0 |
| *variabilis* | 0 | 0 | 0 | 0 | 0 | 0 | 0 | 0 | 0 | 0 | 0 | 1 | 1 | 0 |
| *wasmanni* | 0 | 0 | 0 | 0 | 1 | 0 | 1 | 0 | 1 | 1 | 0 | 0 | 0 | 0 |

**Table S3.** Adjacency matrix on the defined areas in the last 30 Mys. 1 = Areas directly connected; 0 = Areas not connected, or indirectly.

| Time period | Areas | 1 | 2 | 3 | 4 | 5 | 6 | 7 | 8 | 9 | 10 | 11 | 12 | 13 | 14 |
| --- | --- | --- | --- | --- | --- | --- | --- | --- | --- | --- | --- | --- | --- | --- | --- |
| 0 - 10Mya | 1 - Southern Africa | 1 |  |  |  |  |  |  |  |  |  |  |  |  |  |
|  | 2 - Sahelian and Eastern Africa | 1 | 1 |  |  |  |  |  |  |  |  |  |  |  |  |
|  | 3 - North East Africa | 0 | 1 | 1 |  |  |  |  |  |  |  |  |  |  |  |
|  | 4 - North West Africa | 0 | 1 | 1 | 1 |  |  |  |  |  |  |  |  |  |  |
|  | 5 - South West Europe | 0 | 0 | 0 | 0 | 1 |  |  |  |  |  |  |  |  |  |
|  | 6 - North West Europe | 0 | 0 | 0 | 0 | 1 | 1 |  |  |  |  |  |  |  |  |
|  | 7 - Italian Peninsula | 0 | 0 | 0 | 0 | 1 | 1 | 1 |  |  |  |  |  |  |  |
|  | 8 - North East Europe | 0 | 0 | 0 | 0 | 0 | 1 | 0 | 1 |  |  |  |  |  |  |
|  | 9 - Balkans | 0 | 0 | 0 | 0 | 0 | 1 | 1 | 1 | 1 |  |  |  |  |  |
|  | 10 - Türkiye and Caucasus | 0 | 0 | 0 | 0 | 0 | 0 | 0 | 1 | 1 | 1 |  |  |  |  |
|  | 11 - Arabian Peninsula | 0 | 0 | 1 | 0 | 0 | 0 | 0 | 0 | 0 | 1 | 1 |  |  |  |
|  | 12 - Irano-Indian Asia | 0 | 0 | 0 | 0 | 0 | 0 | 0 | 0 | 0 | 1 | 1 | 1 |  |  |
|  | 13 - Central Asia | 0 | 0 | 0 | 0 | 0 | 0 | 0 | 1 | 0 | 0 | 0 | 1 | 1 |  |
|  | 14 - Eastern Asia | 0 | 0 | 0 | 0 | 0 | 0 | 0 | 0 | 0 | 0 | 0 | 1 | 1 | 1 |
| 10 - 20Mya | 1 - Southern Africa | 1 |  |  |  |  |  |  |  |  |  |  |  |  |  |
|  | 2 - Sahelian and Eastern Africa | 1 | 1 |  |  |  |  |  |  |  |  |  |  |  |  |
|  | 3 - North East Africa | 0 | 1 | 1 |  |  |  |  |  |  |  |  |  |  |  |
|  | 4 - North West Africa | 0 | 1 | 1 | 1 |  |  |  |  |  |  |  |  |  |  |
|  | 5 - South West Europe | 0 | 0 | 0 | 1 | 1 |  |  |  |  |  |  |  |  |  |
|  | 6 - North West Europe | 0 | 0 | 0 | 0 | 1 | 1 |  |  |  |  |  |  |  |  |
|  | 7 - Italian Peninsula | 0 | 0 | 0 | 0 | 1 | 1 | 1 |  |  |  |  |  |  |  |
|  | 8 - North East Europe | 0 | 0 | 0 | 0 | 0 | 1 | 0 | 1 |  |  |  |  |  |  |
|  | 9 - Balkans | 0 | 0 | 0 | 0 | 0 | 1 | 1 | 1 | 1 |  |  |  |  |  |
|  | 10 - Türkiye and Caucasus | 0 | 0 | 0 | 0 | 0 | 0 | 0 | 1 | 1 | 1 |  |  |  |  |
|  | 11 - Arabian Peninsula | 0 | 0 | 1 | 0 | 0 | 0 | 0 | 0 | 0 | 1 | 1 |  |  |  |
|  | 12 - Irano-Indian Asia | 0 | 0 | 0 | 0 | 0 | 0 | 0 | 0 | 0 | 1 | 1 | 1 |  |  |
|  | 13 - Central Asia | 0 | 0 | 0 | 0 | 0 | 0 | 0 | 1 | 0 | 0 | 0 | 1 | 1 |  |
|  | 14 - Eastern Asia | 0 | 0 | 0 | 0 | 0 | 0 | 0 | 0 | 0 | 0 | 0 | 1 | 1 | 1 |
| 20 - 30Mya | 1 - Southern Africa | 1 |  |  |  |  |  |  |  |  |  |  |  |  |  |
|  | 2 - Sahelian and Eastern Africa | 1 | 1 |  |  |  |  |  |  |  |  |  |  |  |  |
|  | 3 - North East Africa | 0 | 1 | 1 |  |  |  |  |  |  |  |  |  |  |  |
|  | 4 - North West Africa | 0 | 1 | 1 | 1 |  |  |  |  |  |  |  |  |  |  |
|  | 5 - South West Europe | 0 | 0 | 0 | 0 | 1 |  |  |  |  |  |  |  |  |  |
|  | 6 - North West Europe | 0 | 0 | 0 | 0 | 1 | 1 |  |  |  |  |  |  |  |  |
|  | 7 - Italian Peninsula | 0 | 0 | 0 | 0 | 1 | 0 | 1 |  |  |  |  |  |  |  |
|  | 8 - North East Europe | 0 | 0 | 0 | 0 | 0 | 1 | 0 | 1 |  |  |  |  |  |  |
|  | 9 - Balkans | 0 | 0 | 0 | 0 | 0 | 0 | 1 | 0 | 1 |  |  |  |  |  |
|  | 10 - Türkiye and Caucasus | 0 | 0 | 0 | 0 | 0 | 0 | 0 | 0 | 1 | 1 |  |  |  |  |
|  | 11 - Arabian Peninsula | 0 | 1 | 1 | 0 | 0 | 0 | 0 | 0 | 0 | 1 | 1 |  |  |  |
|  | 12 - Irano-Indian Asia | 0 | 0 | 0 | 0 | 0 | 0 | 0 | 0 | 0 | 1 | 1 | 1 |  |  |
|  | 13 - Central Asia | 0 | 0 | 0 | 0 | 0 | 0 | 0 | 1 | 0 | 0 | 0 | 1 | 1 |  |
|  | 14 - Eastern Asia | 0 | 0 | 0 | 0 | 0 | 0 | 0 | 0 | 0 | 0 | 0 | 1 | 1 | 1 |

**Table S4.** Calibrated nodes before datation inference. Obtained from Branstetter *et al*. (2022).

| Node | Calibration 95%HPD |
| --- | --- |
| Root Stenammini | [42.5482 - 60.1302] |
| [Stenamma] | [29.9377 - 51.2365] |
| [Veromessor] [Novomessor] | [16.1763 - 31.4011] |
| [Veromessor + N] [D + G + O + A + M] | [41.2472 - 58.5796] |
| [Deromyrma] [G + O + A + M] | [33.8794 - 51.5332] |

**Table S5.**
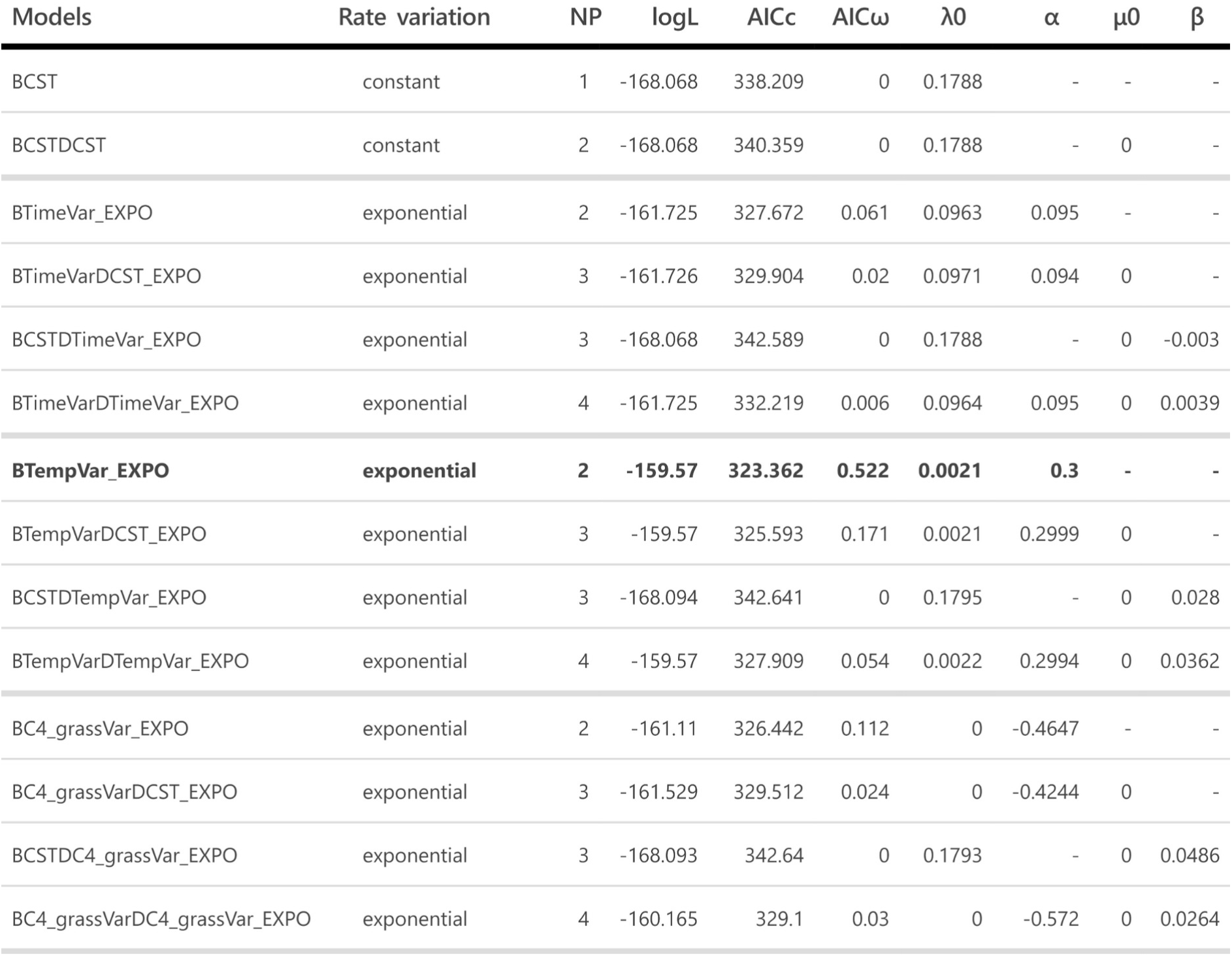
Palaeoenvironment-dependent diversification for the *Messor* ants. The model with temperature dependence speciation and no extinction had the lowest AICc and highest AICω. The analyses show a positive correlation (α>0) for the clade, suggesting higher speciation rates during warm periods. Abbreviations: B, birth (speciation); D, death (extinction); CST, constant rate; Time, time-variable rate; Temp, temperature-variable rate; C4_grass, grassland-variable rate; NP, number of parameters; logL, log-likelihood; AICc, corrected Akaike Information Criterion; AICω, the Akaike weight of each model. Parameter estimates: λ0 and µ0, speciation and extinction for the environmental value at present; α and β, parameters controlling the variation of the speciation (α) and extinction (β) with temperature, C_4_ grasses, or time.

